# Transcriptome-wide association study of schizophrenia and chromatin activity yields mechanistic disease insights

**DOI:** 10.1101/067355

**Authors:** Alexander Gusev, Nick Mancuso, Hilary K Finucane, Yakir Reshef, Lingyun Song, Alexias Safi, Edwin Oh, Schizophrenia Working Group of the Psychiatric Genomics Consortium, Steven McCarroll, Benjamin Neale, Roel Ophoff, Michael C O’Donovan, Nicholas Katsanis, Gregory E Crawford, Patrick F Sullivan, Bogdan Pasaniuc, Alkes L Price

## Abstract

Genome-wide association studies (GWAS) have identified over 100 risk loci for schizophrenia, but the causal mechanisms remain largely unknown. We performed a transcriptome-wide association study (TWAS) integrating expression data from brain, blood, and adipose tissues across 3,693 individuals with schizophrenia GWAS of 79,845 individuals from the Psychiatric Genomics Consortium. We identified 157 genes with a transcriptome-wide significant association, of which 35 did not overlap a known GWAS locus; the largest number involved alternative splicing in brain. 42/157 genes were also associated to specific chromatin phenotypes measured in 121 independent samples (a 4-fold enrichment over background genes). This high-throughput connection of GWAS findings to specific genes, tissues, and regulatory mechanisms is an essential step toward understanding the biology of schizophrenia and moving towards therapeutic interventions.

## INTRODUCTION

Genome-wide association studies (GWAS) have yielded thousands of robustly associated variants for schizophrenia (SCZ) and many other complex traits, but relatively few of these associations have implicated specific biological mechanisms^1,2^, as GWAS association signals often span many putative target genes, may affect gene expression through regulatory^3^ or structural elements^4^, and may affect genes at considerable genomic distances via chromatin looping^5,6^. A growing body of research has demonstrated the enrichment of SCZ GWAS risk variants and heritability within regulatory elements identified through maps of chromatin modifications and accessibility^1,7–13^. Since chromatin modifications are themselves under genetic control^6,14–19^, a causal mechanism for SCZ loci could lead from genetic variation to chromatin modifiers to gene expression and finally to disease risk. Indeed, QTLs for chromatin (and other molecular phenotypes) are enriched within GWAS associations, further supporting this hypothesis^6,18,20,21^.

In this work, we leveraged large gene expression cohorts from multiple tissues, as well as splice variants in brain, to perform a transcriptome-wide association study (TWAS)^22–24^ in a large SCZ GWAS data set^1^ to identify genes whose expression is associated with SCZ and mediated by genetics. We subsequently performed a TWAS for a diverse set of chromatin phenotypes to identify SCZ susceptibility genes that are also associated with specific regulatory elements. To our knowledge, this is the first TWAS to integrate analysis of gene expression, differential splicing, and chromatin variation, moving beyond top SNPs to implicate SCZ-associated molecular features across the regulatory cascade (Figure 1A).

**Figure 1:**
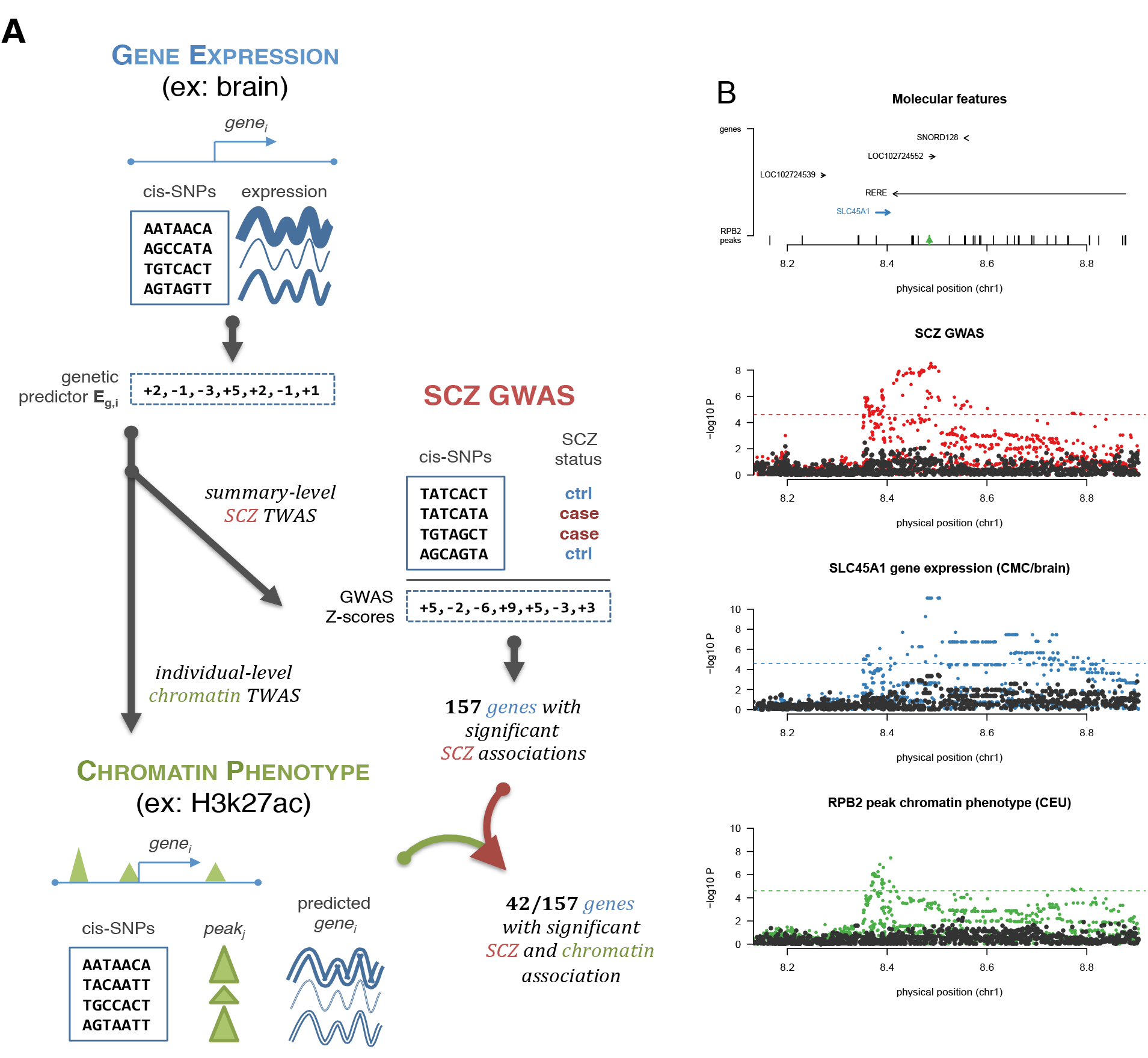
Schematic and example of TWAS approach. (**left**) Illustration of the TWAS approach: genetic predictor of gene expression (*E_g_* is learned in a reference panel (top); integrated with SCZ GWAS association statistics to infer SCZ-*E_g_* association (middle); further integrated with individual-level chromatin phenotypes to infer genes with SCZ and chromatin-¾ associations (bottom), **(right)** Example association of *SLC45Al* gene expression and SCZ, as well as *SLC45Al* expression and a distal RPB2 chromatin peak. Top panel shows locus schematic with all nearby genes and chromatin peaks and corresponding associations highlighted. Three lower panels show Manhattan plots of marginal association statistics before and after conditioning on the TWAS predicted expression (colored/dark dots, respectively). Dashed line shows local significance threshold after Bonferroni correction for number of SNPs. The full 1MB cis locus harbors 15 genes and 60 RPB2 peaks.

## RESULTS

### TWAS for SCZ identifies new susceptibility genes

We analyzed gene expression and genome-wide SNP array data in 3,693 individuals across four expression reference panels spanning three tissues: RNA-seq from the dorsolateral prefrontal cortex (PFC) of 621 individuals - including SCZ and bipolar (BIP) cases and controls - collected by the CommonMind Consortium (CMC)^25^ (see Web Resources), expression array data measured in peripheral blood from 1,245 unrelated individuals from the Netherlands Twin Registry (NTR) ^26^, expression array data measured in blood from 1,264 individuals from the Young Finns Study (YFS) ^23^, and RNA-seq measured in adipose tissue from 563 individuals from the Metabolic Syndrome in Men study (METSIM)^23^. The CMC/brain RNA-seq data further allowed the characterization of differentially spliced introns^27^ (see Methods). Average cis and trans estimates of SNP-heritability of expression (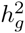, see Methods) were highly significant in each panel, with nominally significant 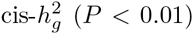 for a total of 18,084 genes summed across the four panels (10,819 unique genes; Table S1), as well as an additional 9,009 differentially spliced introns in brain (in 3,908 unique genes; Table S1).

We performed a TWAS using each of the four gene expression reference panels and summary-level data from the PGC SCZ GWAS of 79,845 individuals^1^ in order to identify genes associated to SCZ (Figure S1). Briefly, this approach integrates information from expression reference panels (SNP-expression correlation), GWAS summary statistics (SNP-SCZ correlation), and LD reference panels (SNP-SNP correlation) to assess the association between the cis-genetic component of expression and phenotype (expression-SCZ correlation)^23^ (Figure 1A). In practice, the expression reference panel was used as the LD reference panel, and cis SNP-expression effect sizes were estimated using a sparse mixed linear model^28^ (see Methods).

The TWAS identified 247 transcriptome-wide significant gene-SCZ and intron-SCZ associations (summed across expression reference panels) for a total of 157 unique genes, including 49 genes that were significant in more than one expression panel (Figure 2, Figure S1, Table 1, Table S2, S3). Of the 104 (non-HLA, autosomal) known PGC GWAS loci^1^, 47 loci overlapped with at least one TWAS gene locus (accounting for 122/157 genes) with the remaining 35/157 genes implicating novel loci. We excluded the MHC region (chr6:28–34MB) from our primary analyses due to its complex haplotype and LD structure. However, as a positive control we specifically tested the *C4A* gene recently fine-mapped for SCZ^4^, which lies inside the MHC, and confirmed a highly significant TWAS association between *C4A* expression in brain tissue and SCZ (P =1.8 × 10^−18^). Across all TWAS associations, the implicated gene was the nearest gene to the top SNP at the locus in only 56% of instances (using the 10,819 cis-heritable genes as background; decreasing to 24% of instances when using all 26,469 known RefSeq genes), underscoring previous observations that the nearest gene to a GWAS hit is often not the most likely susceptibility gene when integrated with expression data^23,24,29,30^. Likewise, conditioning on the predicted expression of a TWAS-associated gene (using summary-level data^31^, see Methods) reduced the *χ*^2^ of the lead GWAS SNP at the locus (including genome-wide significant and non-significant loci) from 42 to 10 on average, and explained more of the association signal than conditioning on the corresponding top expression-QTL (eQTL) (Table S4). For the 43 lead GWAS SNPs at genome-wide significant loci that were in LD (r^2^ > 0.05) with the predicted expression of at least one TWAS-significant gene (out of 47 overlapping index SNPs), joint conditioning on the predicted expression of all such genes reduced the median SNP P-value from *P* =1.2 × 10^−10^ to *P* = 0.028 (Table S5). Given that the TWAS typically captures only 60–80%of the cis component of gene expression at these expression panel sample sizes^23^, the complete elucidation of the cis component could potentially explain the entire GWAS signal at these loci.

**Figure 2:**
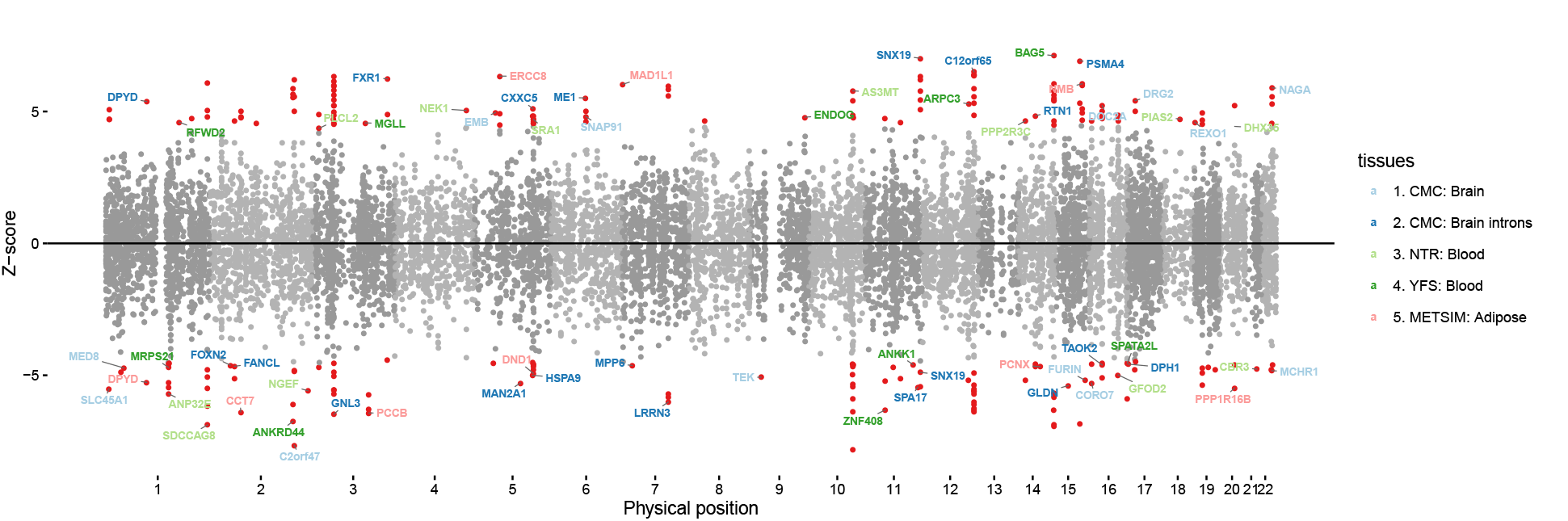
Association between gene expression and schizophrenia. Manhattan plot of SCZ TWAS associations across all tissues. Each point represents a single gene tested, with physical position plotted on x-axis and Z-score of association between gene and SCZ plotted on y-axis. Transcriptome-wide significant associations are highlighted as red points, with “clumped” independent associations (see Methods) labeled with gene names and color-coded by expression reference.

**Table 1:** Number of TWAS-associated genes across all phenotypes and tissues.

|  | CMC/brain introns* | CMC/brain | NTR/blood | YFS/blood | MET/adipose | Total** |
| --- | --- | --- | --- | --- | --- | --- |
| Heritable | (9,009) 3,890 | 5,514 | 2,743 | 5,418 | 4,654 | 11,749 |
| SCZ associated | (80) 46 | 44 | 35 | 48 | 39 | 157 |
| SCZ associated (joint†) | (21) 20 | 13 | 12 | 10 | 9 | 63 |
| SCZ associated (novel) | (12) 10 | 9 | 6 | 6 | 7 | 35 |
| SCZ associated (novel joint†) | (4) 4 | 4 | 3 | 4 | 2 | 17 |
| Chromatin associated | (224) 125 | 244 | 182 | 346 | 232 | 806 |
| SCZ and chromatin associated | (10) 8 | 11 | 10 | 13 | 7 | 42 |
| Enrichment P-value | 8.1e-05 | 2.0e-05 | 4.4e-04 | 6.5e-05 | 2.9e-02 | 1.4e-11 |

\* Number of differentially spliced introns shown in parenthesis.
\*\* Total number of unique gene associations.
† Significant in a summary-based joint analysis.

For both total gene expression and differentially spliced introns, we frequently observed hotspots of multiple TWAS-associated genes in the same locus, a phenomenon previously observed for complex traits^30^. To quantify the total number of independently associated genes, we applied summary statistic-based approximate conditional and joint association methods^31^ to identify genes/introns that had significant TWAS associations when analyzed jointly (see Methods). This yielded a set of 63 jointly significant associations, of which 17 were at novel loci (Table 1, Table S3). 8 loci contained multiple significant associations (spanning 16 genes) in the joint analysis, indicative of allelic heterogeneity. Differentially spliced introns in CMC/brain accounted for more jointly significant associations than any other reference panel (Table 1), followed by gene expression in brain, emphasizing the importance of having both expression and splicing measured in a relevant tissue.

The differentially spliced introns accounted for 46 transcriptome-wide significant gene associations (of which 10 were at novel loci), comparable to the 44 significant gene associations from brain (Table 1, Supplementary Materials), despite the fact that differentially spliced introns accounted for 30% fewer significantly cis-heritable genes than total expression (Table S1). Overall, 20/46 associations corresponded to genes that were not tested in the analysis of total gene expression due to non-significant expression heritability, and 19 of the remaining 26 did not have a transcriptome-wide significant association for total gene expression. We identified multiple TWAS loci driven by specific splice-QTL (sQTL) SNPs that significantly explained a SCZ GWAS association independent of the eQTL effects (Figure S2, S3, S4; Supplementary Materials). This is consistent with the recent observation that sQTLs are typically independent of eQTLs at the same gene^27^. We note that, in contrast to total gene expression, effect direction for these associations is difficult to interpret because multiple alternatively spliced exons within an isoform yielded excised introns that were highly negatively correlated.

This SCZ GWAS data^1^ was recently evaluated in a TWAS with gene expression using Summary-based Mendelian Randomization (SMR) ^24^, identifying 16 transcriptome-wide significant associated genes (in contrast to 157 identified here). This gap could be explained by several differences in the methods: SMR relies on individual eQTL significance and a first stage association at the top eQTL, which has been previously shown to have less power than our TWAS approach using all SNPs in the locus^23^; the SMR test uses an intentionally conservative second stage test for heterogeneity; and the SMR analysis used a different expression panel from a meta-analysis of blood. Of the 16 gene associations identified by SMR, 12 were tested in our study in blood, all replicated at nominal *P* < 0.05 (with consistent sign), and 9 were transcriptome-wide significant - a striking concordance given the different methods and independent expression panels used.

### TWAS associations replicate in internal and external SCZ cohorts

We first replicated the TWAS signal using internal cross-validation within the PGC SCZ GWAS cohorts^1^ (we had permission to access individual genotypes for 58,246 of 79,845 samples). As a prerequisite step, we verified that TWAS using the raw GWAS data produced similar results as TWAS using summary statistics ^23^, observing a correlation of Z-scores ranging from 0.85–0.90 despite the somewhat different set of GWAS samples and independent measures of LD (Table S6). Next, we down-sampled the PGC data into GWAS discovery samples of various sizes (10,000–50,000) to quantify the power and out-of-sample replication of the SCZ TWAS associations (where the expression panel size was always held constant; Figure 3A). We estimated TWAS effect-size precision as the slope of a regression of replication vs. discovery TWAS effect sizes across transcriptome-wide significant associations (see Methods), where a slope below 1 represents over-estimated effect sizes in the discovery data due to winner’s curse. Across random down-samples, average effect-size precision was 0.93 at a discovery sample size of 50,000, indicating minimal winner’s curse and projecting highly accurate effect-size estimates in the full TWAS. The number of transcriptome-wide significant associated genes increased linearly with GWAS sample size and was similar across all four expression reference panels (with more associations in brain expression at the largest discovery size), consistent with a polygenic architecture and many undiscovered associations.

**Figure 3:**
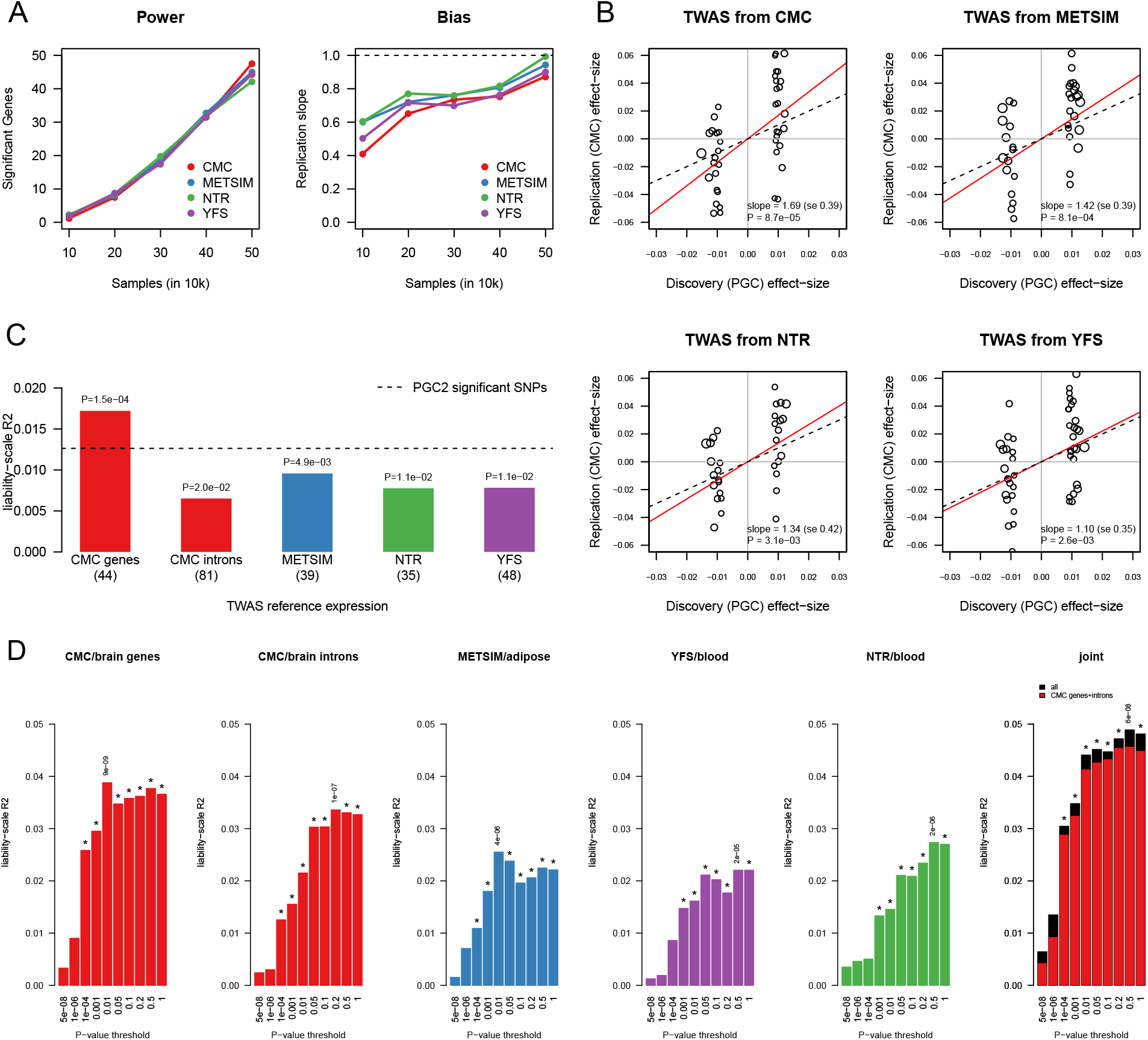
Replication of TWAS associations. (a) The PGC data was randomly split into increasingly large discovery samples (size on the x-axis) and TWAS statistics were estimated from each reference panel. Left panel reports the number of significant genes (after 5% FWER correction) for a given GWAS sample size. Right panel reports the slope from a regression of *β*_replication_ ~ *β*_discovery_ for significant genes identified at each sample size (where all 56, 000 – *x* remaining samples were used as replication). (b) TWAS effectsizes for association to schizophrenia identified in the PGC (x-axis) compared to corresponding estimates in the CMC (y-axis). Dotted line corresponds to *y* = *x*; red line corresponds to the slope from a (Z-score weighted) regression of CMC ~ PGC, with estimate and p-value shown in bottom right. (c) Schizophrenia risk prediction *R*^2^ shown for risk scores constructed from significant TWAS genes (bars) and PGC2 GWAS SNPs (dashed line, comparable to the 1% – 3% reported in ref^1^). Number of genes used in each score reported in parenthesis. Linear-regression *R*^2^ of phenotype on predictor (after subtracting *R*^2^ from jointly fit ancestry PCs) was transformed to the liability scale assuming schizophrenia prevalence of 1%, a linear transformation consistent across all predictors. (d) Schizophrenia risk prediction *R*^2^ for polygenic gene risk scores across multiple significance thresholds. Significant correlations (after Bonferroni correction for number of thresholds tested) are indicated with a (*) and most significant P-value reported. Right-most panel shows prediction from all tissues jointly (black) and from CMC/brain genes + differentially expressed introns jointly (red).

We next replicated the TWAS associations externally using case-control phenotypes from the CMC cohort (which consisted of SCZ+BIP cases and controls; all cases were included due to the high genetic correlation between these two diseases ^32^). We observed significant replication across all four expression panels, with effect-size precision not significantly different from 1 (Figure 3B; Figure S5). Surprisingly, the TWAS gene effect-sizes achieved a higher effect-size precision than GWAS SNP effect-sizes (Figure S6), possibly due to TWAS aggregating heterogeneous effects in a locus more effectively than a single top SNP. We note that even though the same CMC samples were used for the TWAS brain expression reference panel and replication using case-control status, this is an independent replication because CMC case-control status was never used in the discovery TWAS.

### Gene-based risk scores are more predictive than top SNPs

Next, using all transcriptome-wide significant TWAS genes and differentially spliced introns identified in the PGC and their corresponding discovery effect sizes, we constructed gene-based risk scores (GeRS) from their predicted expression in the CMC (SCZ+BIP) case-control samples (see Methods). The GeRS from each expression panel were significantly associated to case-control status, with the strongest association coming from the CMC/brain expression GeRS, which explained 1.4× more SCZ variance (and was more significant; *P* = 2.0 × 10^−4^ vs. 1.2 × 10^−3^) than a genetic risk score computed using the 104 (non-HLA, autosomal) published genome-wide significant GWAS associations^1^ (Figure 3C). The GeRS remained significant in a joint model with the published GWAS predictor (*P* = 0.01, Table S7) showing that the TWAS is prioritizing disease-relevant signal beyond the top SNPs.

We relaxed the significance threshold and found gene-based polygenic risk scores (GePRS) to be highly predictive across the full spectrum of TWAS association P-values (Figure 3D), as observed previously with SNP-based polygenic scores^1,33,34^. Although the prediction was significant in all tissues individually, there was evidence of increased effect in brain, with the prediction from brain (genes and introns) capturing 92% of the joint prediction from all tissues (Figure 3D; Figure S7). A GePRS from actual measured expression and differential splicing in brain was substantially less significant than the genetic GePRS (Figure S7), as expected if the non-genetic component of expression is independent of the TWAS signal. Based on polygenic theory^35,36^, the best TWAS GePRS was estimated to account for 26% of the total SCZ SNP-heritability (see Supplementary Material), indicating a substantial contribution from cis effects in these four tissues.

We sought to investigate temporal differences within the CMC/brain TWAS signal using individual transcriptomes collected by the BRAINSPAN study (see Web Resources) across developmental periods in the same brain sub-region (PFC) ranging from fetal to adult. For each of 19 developmental periods, we estimated differential expression relative to the other periods as an indicator of temporal specificity. We observed TWAS *χ^2^* statistics to be significantly positively correlated with differential expression from the mid-fetal developmental period (*P* < 0.05/19), and corresponding significant negative correlation for differential expression from post-fetal periods (Figure S8, S9). The effect was most significant in the CMC/brain TWAS, less significant in the two blood TWAS, and non-significant in the adipose TWAS. This further underscores tissue-specific differences in the TWAS associations, and prioritizes genes expressed in early development for relevance to SCZ.

### Chromatin TWAS identifies specific regulatory mechanisms for SCZ-associated genes

We next sought to identify relationships between the expression of associated genes from the SCZ TWAS and variation in cis-regulatory elements marked by chromatin (Figure 1A). We used population-level ChIP-Seq chromatin phenotypes measured in 76 HapMap YRI LCLs for H3k27ac (marking active enhancers), H3k4me1 (enhancers), H3k4me3 (promoters), and DNAse (open chromatin)^6^, and in 45 HapMap CEU LCLs for H3k27ac, H3k4me1, H3k4me3, PU1 (regulatory transcription factor) and RNA polymerase II (RPB2, associated with active transcription)^18^. For each of the nine chromatin phenotypes, sites with an excess of ChIP-Seq reads were categorized into local “peaks” corresponding to increased chromatin activity^6,18^. For each peak, the chromatin abundance across individuals was then treated as a single quantitative trait (with quality control mirroring the gene expression analyses; see Methods). Both cohorts additionally had gene expression measured by RNA-seq in the same samples, and we first used Haseman-Elston regression^37^ to quantify the average genetic correlation between gene expression and all chromatin phenotype peaks in the cis locus, for all genes (see Methods). These cis genetic correlations were highly significant (and substantially higher than total correlation of measured phenotypes) across all chromatin phenotypes, persisting for peaks as far as 500kb from the TSS (Figure S10, S11, Table S8, see Methods). We also observed large and highly significant genetic correlations between peaks from different chromatin phenotypes (Figure S12), suggesting that such peaks may tag a single underlying biological feature.

Motivated by these findings, we applied individual-level TWAS methods^23^ to predict expression from the much larger expression reference panels into samples with chromatin phenotypes and searched for expression-chromatin associations. Prediction was performed from expression to chromatin phenotype samples (instead of from chromatin phenotype to expression samples) due to improved numerical stability in the larger expression panels, but we note that this choice was agnostic to the direction of causality (See Supplementary Materials). We confirmed by simulation that this chromatin TWAS strategy is well-calibrated (Figure S13) and much better powered to identify SNP → chromatin → expression associations compared to the conventional approach of testing each SNP for a significant association to both expression (eQTL) and nearby chromatin peaks (cQTL)^6,18^ (Figure 4A, Figure S14).

**Figure 4:**
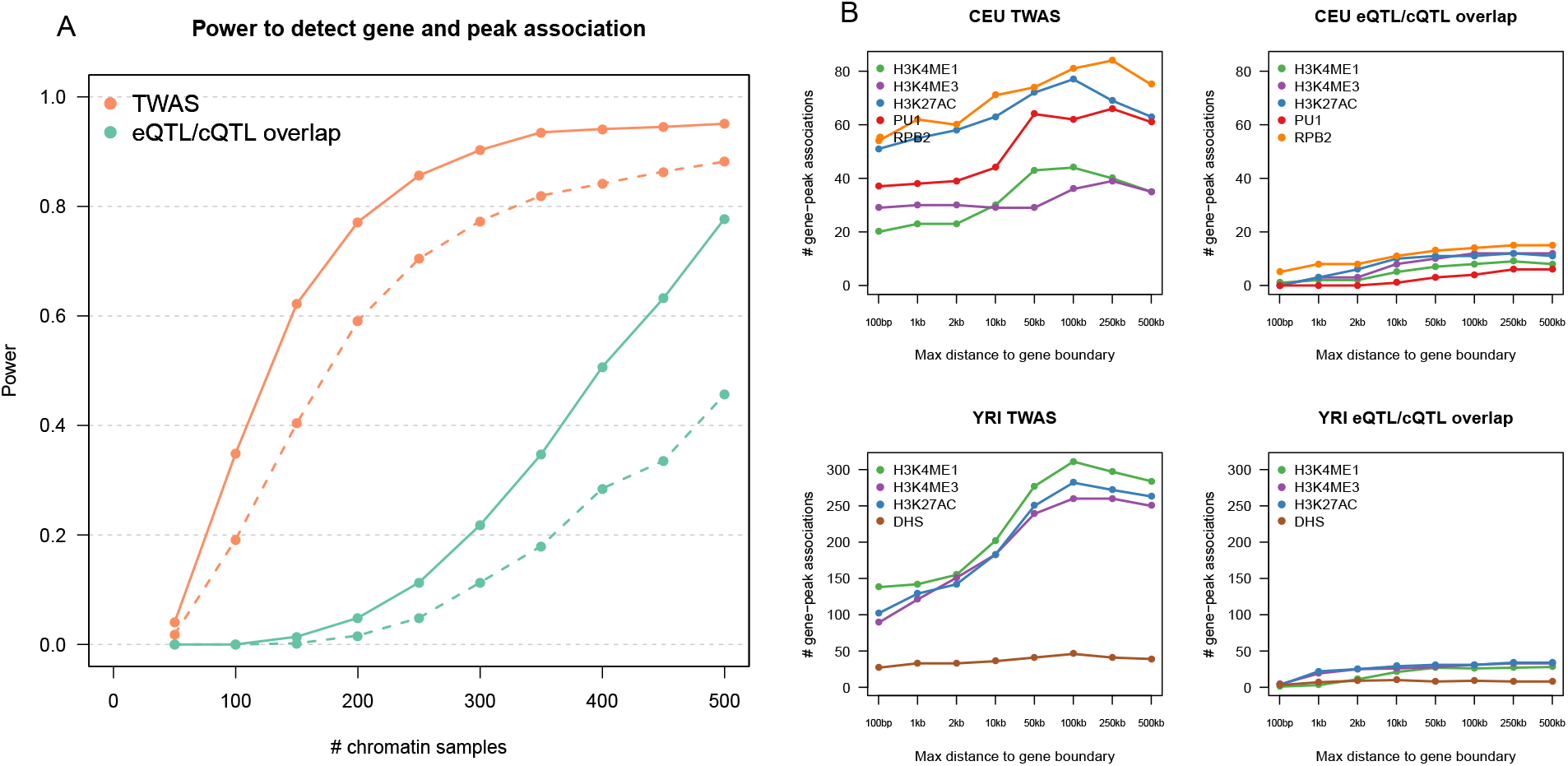
Power and detection of significant gene-mark associations. (a) Molecular phenotypes were simulated under the SNP → chromatin → expression model and two methods to detect gene-mark associations evaluated. (TWAS, orange) corresponds to predicting expression from a held-out reference panel with 1,000 individuals and testing each proximal chromatin peak for association. (eQTL/cQTL, green) corresponds to identifying SNPs that are significantly associated with both chromatin and expression at the locus. For a given chromatin phenotype sample size (x-axis), power was measured as the number of instances where locus was deemed significant after accounting for number of gene-mark pairs tested (TWAS) or number of SNPs and gene-mark pairs tested (eQTL/cQTL). Solid (dashed) lines correspond to 1 (2) chromatin-causing variants in the simulation. (b) Genes significantly associated with a chromatin phenotype peak in YRI/CEU. For a given distance from the gene (x-axis), the number of unique genes is reported after experiment-wide Bonferroni correction for tests across all chromatin phenotypes. Results from TWAS prediction-based approach shown on left; results from overlapping QTL approach on the right (see Methods).

We performed the chromatin TWAS for the 10,819 significantly heritable genes and 9,009 differentially spliced introns analyzed in the SCZ TWAS, together with all chromatin peaks in the ±500kb locus of each gene. Focusing on the 157 transcriptome-wide significant genes from the SCZ TWAS, we identified 42 genes (including 7 genes at novel loci) that also had Bonferroni significant chromatin TWAS associations (to a total of 78 individual chromatin peaks) in analyses using the same expression reference panel (Table 1, Table 2, Table S9, S10). Significant evidence of a chromatin-SCZ association was observed for the majority of genes using a separate TWAS-like test as well as the SMR^24^ test using cQTLs, in spite of the low sample size (Supplementary Materials, Table S11). Individually, these 42 loci represent specific mechanistic hypotheses where the implicated chromatin phenotypes are disrupted and mediate the expression of susceptibility genes. Overall, there was a highly significant enrichment of gene-chromatin TWAS associations at SCZ TWAS genes relative to all heritable genes (OR=3.9; *P* =1.4 × 10^−11^ by Fisher’s exact test), suggesting that such mechanisms are particularly relevant for SCZ susceptibility genes. The enrichment was also individually significant (*P* < 0.05/5) for all tissues except adipose (Table 1). Surprisingly, the SCZ-associated differentially spliced introns were also enriched for chromatin associations (0R=5.0, *P* = 8.1 × 10^−5^), even though differentially spliced introns had many fewer chromatin associations overall (see below), suggesting that epigenetic regulation of splicing may be particularly relevant for SCZ susceptibility genes. Only 8 of the 78 chromatin peaks underlying joint SCZ TWAS and chromatin TWAS associations were within the promoter (±2kb of the TSS) of their associated gene. This suggests that most regulatory elements affecting SCZ are distally located, as previously observed in other traits^6,8,20^ and underscores the importance of searching broadly around the TSS.

**Table 2:** TWAS genes with association to schizophrenia and chromatin phenotypes. 42 genes (including 7 genes at novel loci, highlighted with a [*]) had a significant TWAS association with SCZ and chromatin phenotypes. For each significant TWAS association with SCZ, the number of significant gene-chromatin associations (FWER 5% among TWAS gene-mark associations, by Bonforroni correction) are reported. In the middle columns ‘.’ represents genes that were not heritable in the study and therefore not TWAS-associated. In the right columns ‘.’ represents no identified association; genes with no chromatin associations are not shown. Top panel shows results from genes, with TSS listed as position; bottom panel shows results from differentially spliced introns in CMC with exon-exon junction listed as position (details in Table S9).

| Gene | Chr | Position | TWAS P-value |  |  |  | Associated marks |  |  |  |  |  |
| --- | --- | --- | --- | --- | --- | --- | --- | --- | --- | --- | --- | --- |
|  |  |  | YFS/blood | MET/adipose | NTR/blood | CMC/brain | DHS | H3K27AC | H3K4ME1 | H3K4ME3 | PU1 | RPB2 |
| RERE | 1 | 8,483,747 | 4e-07 | 2e-06 | 2e-06 | . | . | . | . | . | . | 3 |
| SLC45A1 | 1 | 8,378,144 | . | . | . | 4e-08 | . | . | . | . | . | 1 |
| MAP7D1* | 1 | 36,621,565 | 6e-04 | . | 1e-06 | . | . | . | . | . | . | 1 |
| MED8 | 1 | 43,855,483 | 5e-01 | . | . | 2e-06 | . | . | 1 | . | . | . |
| ANP32E | 1 | 150,207,026 | . | . | 1e-08 | . | . | . | . | 1 | . | . |
| MRPS21 | 1 | 150,266,261 | 3e-06 | 3e-03 | 6e-03 | 2e-02 | . | . | 1 | . | . | . |
| RFWD2* | 1 | 176,176,380 | 4e-06 | . | . | . | . | . | 1 | . | . | . |
| C2orf69 | 2 | 200,775,978 | . | 6e-10 | . | . | . | . | . | 1 | . | . |
| GLT8D1 | 3 | 52,737,714 | . | . | 5e-08 | 3e-08 | . | 1 | . | . | . | . |
| GLYCTK | 3 | 52,321,835 | 2e-08 | . | . | . | . | 1 | . | . | . | . |
| GNL3 | 3 | 52,719,935 | 7e-09 | 6e-07 | . | 5e-02 | . | . | . | 1 | . | . |
| NEK4 | 3 | 52,804,965 | . | . | . | 2e-09 | . | . | . | 1 | . | . |
| NT5DC2 | 3 | 52,567,793 | 6e-06 | 6e-06 | . | 7e-01 | . | 1 | . | 1 | . | . |
| PPM1M | 3 | 52,279,808 | 2e-07 | 2e-07 | . | 2e-03 | . | 1 | . | . | . | . |
| TMEM110 | 3 | 52,931,597 | 1e-02 | 4e-01 | 1e-08 | 6e-06 | . | 1 | 1 | 2 | . | . |
| PCCB | 3 | 135,969,166 | 1e-08 | 1e-10 | . | 3e-10 | 1 | . | 3 | . | . | . |
| RP11-53O19.3 | 5 | 44,826,178 | . | 6e-06 | . | . | . | . | 1 | . | . | . |
| DND1 | 5 | 140,053,171 | . | 8e-07 | 1e-02 | . | . | 1 | . | 1 | . | . |
| IK | 5 | 140,027,383 | 4e-06 | 1e-06 | . | 5e-05 | . | 1 | . | 1 | . | . |
| NDUFA2 | 5 | 140,027,370 | 2e-06 | . | . | 4e-06 | . | 1 | . | 2 | . | . |
| PCDHA2 | 5 | 140,174,443 | . | . | . | 7e-06 | . | 1 | . | 1 | . | . |
| ZMAT2 | 5 | 140,080,031 | 5e-06 | 1e-03 | . | 3e-06 | . | . | . | 1 | . | . |
| AS3MT | 10 | 104,629,209 | . | 6e-08 | 7e-09 | 1e-05 | . | 1 | . | . | . | . |
| MPHOSPH9 | 12 | 123,717,785 | 4e-09 | 1e-05 | . | 2e-08 | . | 1 | . | . | 1 | . |
| KIAA0391* | 14 | 35,591,526 | 7e-01 | 2e-07 | 5e-01 | . | . | 2 | 1 | . | . | . |
| PPP2R3C* | 14 | 35,591,748 | 6e-05 | 1e-01 | 3e-06 | 2e-02 | . | 2 | 2 | . | . | . |
| MAPK3 | 16 | 30,134,630 | 5e-05 | . | . | 1e-06 | . | 1 | . | . | . | 1 |
| GFOD2 | 16 | 67,753,273 | . | . | 6e-07 | 2e-05 | . | 1 | . | 2 | . | . |
| TSNAXIP1 | 16 | 67,840,780 | . | . | . | 2e-06 | . | . | 1 | 2 | . | . |
| DUS2L | 16 | 68,038,024 | 1e-06 | . | 3e-06 | 4e-04 | . | . | . | 2 | . | . |
| PRMT7 | 16 | 68,344,876 | 1e-05 | 8e-04 | . | 8e-06 | . | . | 1 | 1 | . | . |
| GRAP* | 17 | 18,950,336 | . | . | 5e-07 | . | . | . | . | . | . | 1 |
| RNF112* | 17 | 19,314,490 | 8e-06 | . | . | . | . | . | . | . | . | 1 |
| ACTR5 | 20 | 37,377,096 | 2e-07 | 2e-04 | . | 7e-01 | 1 | . | 1 | . | . | . |
| CBR3 | 21 | 37,507,262 | 6e-03 | 2e-03 | 2e-06 | 5e-04 | 1 | . | 2 | . | . | . |
| CMC/brain splicing |  |  |  |  |  |  |  |  |  |  |  |  |
| TBC1D5 | 3 | 17,255,862 - 17,279,655 | . | . | . | 3e-06 | . | . | 1 | . | . | . |
| NEK4 | 3 | 52,800,010 - 52,800,194 | . | . | . | 1e-06 | . | . | 1 | . | . | . |
| CCDC90B | 11 | 82,985,783 - 82,991,184 | . | . | . | 3e-07 | . | 1 | . | . | . | . |
| SBNO1 | 12 | 123,821,038 - 123,825,535 | . | . | . | 4e-10 | . | . | . | . | 1 | . |
| KLC1 | 14 | 104,145,855 - 104,151,323 | . | . | . | 7e-12 | . | . | 1 | 1 | . | . |
| RTN1* | 14 | 60,074,210 - 60,193,637 | . | . | . | 1e-06 | . | . | 1 | . | . | . |
| TAOK2 | 16 | 29,997,825 - 29,998,165 | . | . | . | 4e-06 | . | . | . | . | . | 1 |
| PPP4C | 16 | 30,094,168 - 30,094,715 | . | . | . | 2e-06 | . | . | . | . | . | 1 |

\* Novel, not overlapping 108 PGC SCZ GWAS loci

We describe three specific examples of TWAS associations to SCZ and chromatin phenotypes. First, the expression of *SLC45A1* in CMC/brain was associated with SCZ (*P* = 3.5 × 10^−8^; overlapping a significant GWAS locus) as well as a distal RPB2 peak (106kb from TSS; *P* = 1.5 × 10^−5^), with significant marginal associations (QTLs) for both expression and chromatin (Figure 1B). TWAS prioritized this gene-peak combination from 15 genes and 60 peaks in the 1MB locus. Conditioning^31^ on the predicted expression of *SLC45A1* explained all significant eQTLs/cQTLs and GWAS SNPs in the locus. Notably, *SLC45A1* was not the nearest gene to the top GWAS SNP nor to the associated chromatin peak, and the lead GWAS SNP was only nominally associated with the RPB2 peak. This highlights a chromatin association that could not have been identified using conventional approaches based on proximity or overlapping top hits. Second, the expression of *PPP2R3C* in NTR/blood was associated with SCZ (*P* = 3.4 × 10^−6^) - despite no genome-wide significant SNPs at the locus - as well as two distal peaks for H3k4me1 (minimum *P* = 1.0 × 10^−9^) and two distal peaks for H3k27ac (minimum *P* = 4.1 × 10^−6^) (Figure S15). The four chromatin peaks clustered together physically and suggest a coordinated regulatory effect on expression. Conditioning on the predicted expression of *PPP2R3C* again explained all significant marginal associations for the implicated phenotypes (Figure S15). *PPP2R3C* was the nearest gene to the most significantly associated SNP at the locus and to the implicated chromatin peaks. However, because the locus was not genome-wide significant, this association would not have been identified in a conventional analysis of known GWAS loci. *PPP2R3C* was also recently identified by SMR analysis of SCZ in an independent expression panel^24^; our findings identify specific chromatin features for experimental follow-up. Third, we describe the *MAPK3* locus in detail below.

### Allelic imbalance analyses localize TWAS association for MAPK3

We highlight the SCZ TWAS association of *MAPK3* expression in CMC/brain (*P* =1.3 × 10^−06^), overlapping a significant SCZ GWAS locus. Notably, *MAPK3* is located inside the 16p11 copy number variant that has been associated with SCZ and autism^38–41^, and has recently been linked with differential protein abundance in SCZ^42^ and shown to respond to pharmacological targeting in cultured neurons^43^. In our analysis, *MAPK3* was also nominally differentially expressed in the independent CMC (SCZ+BIP) case-control samples (Wilcoxon *P* = 0.03, over-expressed in controls) despite the small sample size. The chromatin TWAS identified associations between *MAPK3* and two peaks near the TSS (H3k27ac, *P* = 7 × 10^−6^; RPB2, *P* =1 × 10^−11^). In the CEU chromatin phenotype samples, where *MAPK3* expression was also measured in LCLs, the H3k27ac and RPB2 peaks explained 36% (*P* = 7 × 10^−6^) and 23% (*P* = 5 × 10^−4^) of the variance in measured expression, respectively, with only the H3k27ac peak significant in a joint model. Because the chromatin TWAS associations were identified in LCLs, we examined additional epigenetic data from H3k27ac, H3k4me3 and ATAC-seq measured in brain tissues (including pre-frontal cortex) as part of the PsychENCODE project^44^ and showed the presence of peaks across all three chromatin phenotypes (Figure S16, Supplementary Materials). Strikingly, both peaks were also nearly identical to two recently identified human-gained neuro-developmental enhancers in independent fetal cortex tissues^45^ (Figure S16), providing compelling evidence of evolutionary importance. We then focused on two putatively functional SNPs within the ATAC-seq peak (rs28529403 and rs61764202, in tight LD with *ρ* = 0.7) that were both recently classified as disrupting multiple transcription factor binding sites^21^ and therefore plausible causal variants. Conditioning on either SNP accounted for all significant marginal associations across the corresponding expression and chromatin phenotypes (Figure S17, S18), consistent with these SNPs driving the local signal.

We sought to confirm the effect at these SNPs by looking at allelic imbalance, which naturally accounts for environmental differences across individuals^46,47^. First, we focused on allele-specific expression in the CMC RNA-seq data (which is statistically independent of the QTL-based signal used for TWAS). We phased the locus and, for each SNP, tested all heterozygous carriers for an imbalance in RNA-seq reads at the transcript SNP (see Methods). Both putative SNPs showed significant evidence of allele-specific expression (rs28529403, *P* = 2.5 × 10^−21^; rs61764202, *P* = 2.0 × 10^−21^), and were the most significantly imbalanced in the locus (Table S12, Figure S19).

Next, we collected additional ATAC-seq data at this locus, for a total of 314 CMC individuals^44^, and looked for allele-specific chromatin activity in the same manner. We restricted to 267 SNPs that were Bonferroni significant for allele-specific expression (above), and tested all heterozygous carriers for an imbalance in ATAC-seq reads using either of the two peak SNPs as anchors (see Methods). We observed a Bonferroni significant allelic imbalance at rs61764202 (94 heterozygous samples, *P* = 8.6 × 10^−5^), which was more significant than any other SNP in phase with rs61764202; we did not observe significant imbalance at rs28529403 (100 samples, *P* = 0.15) or any other SNP in phase with rs28529403 (Table S12). The imbalance was also individually significant in four samples but with opposite direction in one of the four (Figure S20, Table S13). For this sample, we cannot decisively rule out the possibility of artifact, a mis-imputed heterozygous call, confounding from an interaction involving nearby features, or tissue sub-type heterogeneity. However, such biases would not be expected to inflate the combined test across all individuals. Taken together, the allele-specific signal from both molecular phenotypes nominates the alternative allele at rs61764202 as disrupting chromatin activity, increasing *MAPK3* expression, which in turn decreases SCZ risk.

### Chromatin TWAS associations explain the bulk of cis expression regulation

We expanded the chromatin TWAS to all 10,819 heritable genes (not just SCZ-associated genes) in order to evaluate the properties of all chromatin TWAS associations. This yielded roughly 7× more Bonferroni significant expression-chromatin associations than using the conventional in-sample eQTL/cQTL overlap approach^6,18^ (Figure 4B, Table S14), consistent with our previous simulations. Across all tissues, 806 unique genes had a transcriptome-wide significant association (see Methods) with at least one chromatin phenotype (Figure 4B, Table S15), and 4,294 genes were significant at the 10% (per-phenotype) FDR used in previous studies^6,18^ (Table S16). In contrast, only 224 of 9,009 differentially spliced introns in the CMC has a transcriptome-wide significant association with a nearby chromatin mark, corresponding to 2–3 × fewer associations than identified using total CMC gene expression (depending on the chromatin phenotype, Table S17). As expected, the distribution of associated chromatin peaks was centered at the TSS of the corresponding gene (Figure S21). However, additional associations were consistently observed when testing distal peaks (> 10kb) after correcting for the additional tests performed, in contrast to the results from eQTL/cQTL overlap (Figure 4B, S22, S23). We evaluated these distal chromatin TWAS associations against genome-wide Hi-C measured in a reference LCL ^6^, reasoning that truly interacting gene-peak pairs are likely to be in 3-dimensional chromatin contact. Across all chromatin phenotypes, we found distal (≥ 10kb) gene-peak associations to be significantly enriched for Hi-C inferred loops relative to random (distance-matched) gene-peak pairs, with an odds ratio of 2.4 (95% CI 2.2–2.5; *P* < 1 × 10^−16^) for pairs up to 500kb apart (Figure S24).

Our primary analyses leveraged external expression data in larger sample sizes, but we also analyzed directly measured expression in chromatin samples for validation. First, we confirmed that the predicted expression was significantly correlated with measured expression in the chromatin samples (Table S18). Even though the four expression reference panels all contained samples of European ancestry, this correlation was significant for both CEU and YRI target samples; with average ñ^2^ for YRI half as large as for CEU, providing an estimate of the power loss due to differences in linkage disequilibrium (LD) patterns across populations^48^. Next, for every TWAS-associated gene-chromatin peak pair, we measured the association between actual measured expression and the corresponding chromatin phenotype. If the chromatin TWAS association reflects true genetic correlation, we would expect the measured expression to also be highly correlated (barring tissue-specific differences between the expression panels). Indeed, across the 806 chromatin TWAS-associated genes, the correlation between measured expression and an associated chromatin phenotype was highly significant when compared against a distance-matched background null (Figure S10B). In CEU, where cross-population LD differences are minimized, the average individual TWAS-associated chromatin peak explained 21% of the variance in measured expression of its target gene for peaks within 2kb of the TSS, with the subset of distal peaks (2kb-500kb to the gene boundary, 32% of the peaks) explaining 18% (Figure S25, S26, S27, S28). This strikingly high variance explained is comparable to the total cis-hg of these genes (in blood, the tissue type most relevant to LCL; Table S19), implying that the cis-genetic effect on expression may be fully explained by individual chromatin TWAS peaks. We stress that the measured expression in chromatin samples was completely independent of the external expression data used in the chromatin TWAS, ensuring that these estimates were not biased by over-fitting or winner’s curse. For the three chromatin phenotypes that were measured in both CEU and YRI, associated peaks identified in one population were still predictive of association with measured expression in the other (Figure S29, S30, Table S20), lending further support to previous observations that regulatory activity and the resulting impact on disease is stable across ethnicities^16,49^ and supporting our use of measurements from multiple populations.

### Chromatin modifications mediate SNP-expression associations that impact SCZ

We next evaluated whether the SCZ and chromatin TWAS associated loci were more consistent with a chromatin mediating (SNP → chromatin → expression) or expression mediating (SNP → expression → chromatin) causal model leading to SCZ susceptibility. We derived a statistic based on the ratio of cis-genetic covariance (*cov_g_*) between SCZ and the two molecular phenotypes (see Methods). Conceptually, the genetic effect of a given molecular phenotype on SCZ will be attenuated by environmental noise, which will manifest itself as lower *cov_g_* to SCZ for phenotypes further along the molecular cascade. We estimated expression-SCZ and chromatin-SCZ *cov_g_* in the CEU and YRI samples with both chromatin and expression data, using cross-trait LD score regression^50^. Averaging across the 42 chromatin TWAS associations at genes identified in the SCZ TWAS, both the expression-SCZ and chromatin-SCZ *cov_g_* were significantly higher than that of a random background of gene-peak pairs less than 500kb apart, with chromatin-SCZ *cov_g_* 2.5× greater on average than expression-SCZ *cov_g_* and more significantly different from the background (Figure S31, see Methods). This corresponds to the chromatin phenotype explaining 35% of the variance in expression under the model where it is the mediator, with the rest due to environmental or trans variance independent of disease (see Methods). Furthermore, regressing the chromatin phenotype out of expression led to non-significant estimates of expression-SCZ *cov_g_*, but regressing the measured expression out of the chromatin phenotype did not significantly affect the chromatin-SCZ *cov_g_* estimates (Figure S31B). Both observations strongly support a chromatin mediating model (SNP → chromatin → expression) previously hypothesized^15–17^, and extend it to SCZ etiology.

## DISCUSSION

The landmark PGC SCZ GWAS paper stated that “if most risk variants are regulatory, available eQTL catalogues do not yet provide power, cellular specificity, or developmental diversity to provide clear mechanistic hypotheses for follow-up experiments” ^1^. In this work, we apply cutting-edge methodology to data from expression and chromatin activity to provide mechanistic hypotheses. Applying the TWAS approach to SCZ, we identified 157 unique genes with transcriptome-wide significant associations, whose predicted expression explained the bulk of the corresponding GWAS SNP association. Genes below the transcriptome-wide significance threshold continued to be strongly associated with SCZ and exhibited clear preference for expression and splicing in the brain (though this can also reflect expression data quality). Indeed, alternative splicing in the brain yielded the greatest number of independent TWAS associations, highlighting an important source of disease-relevant variation^27^ with potential therapeutic implications^51,52^. 42 of the SCZ-associated genes were significantly associated with nearby chromatin variation (a 4-fold enrichment relative to non-SCZ genes), implicating specific regulatory features for functional follow-up. Our analyses strongly supported a model where chromatin variation (likely marking differences in transcription-factor binding) mediates the relationship between genetics and expression^15–17^, and connect this model with SCZ etiology. This may explain the modest overlap between eQTLs and GWAS hits previously reported^29,30,46^, where the downstream nature of the expression phenotype (relative to chromatin) would decrease power for such overlap analyses.

We note several limitations and future directions of this study. First, the discovery chromatin phenotypes analyzed here were measured in LCLs, preventing us from identifying regulatory elements that were brain-specific and of potentially greater relevance to SCZ. Second, although TWAS is not confounded by reverse-causality (disease → expression), instances where a SNP influences SCZ which in turn influences gene expression (or the same SNP influences SCZ and expression independently) are statistically indistinguishable from causal susceptibility genes. Therefore, although we can conclude that 26% of the SCZ 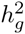 is explained by predicted expression of all genes analyzed here we cannot estimate the fraction that is truly causal without further mediation experiments. Finally, our simulations indicate that the chromatin TWAS approach will have substantially greater power as chromatin sample sizes increase into the hundreds^44^. As tissue acquisition may be the biggest hurdle for producing larger data sets, methods that do not depend on measurements from the same samples will remain critical. Above and beyond specific mechanistic findings for SCZ, our findings outline a systematic approach to identify biological mediators of complex disease.

## WEB RESOURCES

TWAS results:

~~~
http://sashagusev.github.io/chromatinTWAS/
TWAS methods:
http://bogdan.bioinformatics.ucla.edu/software/twas/
BRAINSPAN transcriptomes:
http://www.brainspan.org/static/download.html
CommonMind consortium:
https://www.synapse.org/cmc
Grubert et al data:
http://chromovar3d.stanford.edu
PGC summary data:
https://www.med.unc.edu/pgc/downloads
PLINK:
https://www.cog-genomics.org/plink2
PsychENCODE knowledge portal:
https://www.synapse.org/#!Synapse:syn4921369/wiki/235539
SNPWeights for principal component analysis:
http://www.hsph.harvard.edu/alkes-price/software/
Waszak et al data (provided by the authors):
http://gardeux-vincent.eu/Cell2015/description.peaks.zip,
http://gardeux-vincent.eu/Cell2015/quantified.peaks.zip,
http://gardeux-vincent.eu/Cell2015/quantified.peaks.PEER.centered.zip
~~~

## ACKNOWLEDGEMENTS

We would like to acknowledge Michael Gandal, Bryce van de Geijn, Arthur Ko, Po-Ru Loh, Luke O’Connor, and Paivi Pajukanta for helpful discussions. This research was funded by NIH grants F32 GM106584, R01 GM105857, R01 MH109978 and R01 MH107649. HKF was supported by the Fannie and John Hertz Foundation. We are grateful to the CommonMind Consortium and the PsychENCODE Consortium for making data publicly and readily available.

CMC: Data were generated as part of the CommonMind Consortium supported by funding from Takeda Pharmaceuticals Company Limited, F. Hoffman-La Roche Ltd and NIH grants R01MH085542, R01MH093725, P50MH066392, P50MH080405, R01MH097276, RO1-MH-075916, P50M096891, P50MH084053S1, R37MH057881 and R37MH057881S1, HHSN271201300031C, AG02219, AG05138 and MH06692. Brain tissue for the study was obtained from the following brain bank collections: the Mount Sinai NIH Brain and Tissue Repository, the University of Pennsylvania Alzheimers Disease Core Center, the University of Pittsburgh NeuroBioBank and Brain and Tissue Repositories and the NIMH Human Brain Collection Core. CMC Leadership: Pamela Sklar, Joseph Buxbaum (Icahn School of Medicine at Mount Sinai), Bernie Devlin, David Lewis (University of Pittsburgh), Raquel Gur, Chang-Gyu Hahn (University of Pennsylvania), Keisuke Hirai, Hiroyoshi Toyoshiba (Takeda Pharmaceuticals Company Limited), Enrico Domenici, Laurent Essioux (F. Hoffman-La Roche Ltd), Lara Mangravite, Mette Peters (Sage Bionetworks), Thomas Lehner, Barbara Lipska (NIMH).

The data reported in this paper are tabulated in the Supplementary Materials and archived at: http://sashagusev.github.io/chromatinTWAS/

## METHODS

### Data and quality control

Genotypes and expression from the NTR^26^, YFS^23^, and METSIM^23^ were processed as described in ref^23^ and the corresponding expression weights were downloaded directly from the TWAS web-site (see Web Resources). Genotypes and expression data from the CMC^25^ were processed using the GTEx Consortium guidelines for eQTL analysis of RNA-seq data. Specifically, RNA-seq RPKM was quantile normalized across samples; genes having > 10 individuals with zero reads were removed; each gene was rank-normalized; 15 PEER factors were computed; and the residual expression used.

We used the LeafCutter algorithm^27,53^ to quantify de novo intron excision in the CMC RNA-seq data by clustering reads that spanned intron junctions. These clusters correspond to individual isoforms and enable an estimate of differential intron splicing computed from the ratio of reads spanning an intron relative to the total isoform read count. Splice variants were quantified using LeafCutter default parameters: a minimum of 50 reads per cluster, and a maximum intron length of 500kb. Based on the guidelines in ref. ^53^, the following quality controls were applied to the inferred isoform clusters: clusters having > 10 individuals with zero reads were removed; clusters with < 100 individuals having > 20 reads were removed; and introns with < 5 individuals having non-zero counts were removed. The inferred per-sample abundance for each intron was then treated as a molecular phenotype, normalized, and PEER-corrected as with total expression above. This process identified 123,480 differentially spliced introns, of which 99,562 mapped to canonical gene introns. We treated the differential splicing of these 99,562 introns as quantitative traits in the same manner as total expression.

For genotype data, individuals failing a sex check or having 5% missing SNPs were removed. Additionally, SNPs were removed if they had > 5% missing calls; *P* < 0.05 case-control missing association; *P* < 5 × 10^−6^ Hardy-Weinberg disequilibrium; *P* < 5 × 10^−3^ association to batch; *P* < 5 × 10^−8^ missing haplotype association; or frequency < 1%. Principal components (PCs) were computed using all samples for the NTR, YFS, and METSIM data directly and using SNPweights (v2.1)^54^ for the CMC data, outliers were removed (samples > 6 standard deviations away the mean along any top component), and PCs included as fixed-effects in estimating 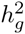. For all datasets, related individuals with GRM values > 0.05 were also removed prior to estimating 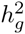.

### 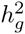 estimation

Cis and trans 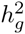 were estimated using variance-components, modeling the phenotype as a multi-variante Normal 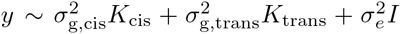 where *K* are the standard genetic relatedness matrices from SNPs in the cis locus (*K*_cis_) and in the rest of the genome (*K*_trans_). The *σ*^2^ parameters were fit for each gene using AI-REML as implemented in the GCTA software^55^, with principal components and sex included as fixed effects. For 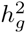 of differentially spliced introns, the intron ratios condition out isoform abundance but total gene expression was also included as a covariate to account for any residual correlation. As in previous studies ^26^, individual estimates outside the plausible 0–1 range were allowed in order to achieve unbiased mean estimates. The standard error of each estimate was approximated as the standard deviation divided by the square root of the number of genes tested; however, significant differences were confirmed by permutation tests (see below).

To evaluate the contribution of low-frequency variants, we imputed the NTR data to the Haplotype Reference Consortium reference, yielding high-quality imputed SNPs down to MAF of 0.001. On average, we did not observe a significantly non-zero contribution of imputed rare variants to 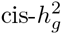, nor did we see a significant change in common 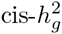 due to denser imputation relative to array SNPs (Table S1). Though recent work has identified biases in estimates of 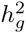 from rare variants^56^, we expect these biases to be small in the cis region and largely mitigated by the two-component model. We did not further evaluate the contribution of rare variants to 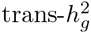. No difference was observed when using dosages to construct the cis GRM.

In the CMC data, where SCZ/BIP and control status was also available, the average cis-genetic correlation of expression between (SCZ/BIP) cases and controls was 1.00 (s.e. 0.02), indicating consistent direction of eQTL effect sizes between cases and controls and motivating us to use the full cohort as a TWAS reference panel (Table S21). We additionally performed multiple analyses of expression heritability associated with functional category (Supplementary Materials) demonstrating pervasive enrichment of chromatin marks near significantly heritable genes and underscoring the importance of chromatin variation to expression heritability ^57^.

### Schizophrenia TWAS

We analyzed publicly available schizophrenia summary statistics from the PGC GWAS of 79,845 individuals^1^ (see Web Resources). Summary statistic-based TWAS was performed as described previously^23^ using cis SNP-expression effect sizes computed by the BSLMM software^28^ for each of the four expression reference panels. Strand-ambiguous alleles (A/T, G/C) were removed from the summary data and all SNPs were set to the same strand. We evaluated TWAS predictions using either the SNPs genotyped in each expression reference panel, or imputed HapMap3 SNPs (which typically represent well-imputed SNPs). To account for multiple hypotheses, we applied Bonferroni correction within each expression panel that was used. This threshold was chosen so as to maximize consistency with previous published results and not penalize for additional (and often highly correlated) expression panels tested. Specifically, we report “transcriptome-wide” significance after correcting for the number of genes tested within each of the five reference panels (CMC, CMC-splicing, NTR, YFS, METSIM; 5,419 tests on average). This is consistent with the correction applied in previous TWAS results of multiple expression references^23^.

To ensure that results from the CMC data were not biased by ascertainment for SCZ/BIP cases, we performed a separate TWAS in which cis SNP-expression effect sizes were stratified on CMC case-control status and meta-analyzed; we observed no significant differences compared to using the full sample (Table S22).

Conditional and joint analysis was performed using the summary statistic-based method described in ref. ^31^, which we applied to genes instead of SNPs. This approach requires marginal association statistics (i.e. the main TWAS results) and a correlation/LD matrix to evaluate the joint/conditional model. The correlation matrix was estimated by predicting the cis-genetic component of expression for each TWAS gene using the CMC genotypes and computing Pearson correlations across all pairs of genes. The 247 transcriptome-wide significant TWAS associations across four reference panels (spanning 157 unique genes) were then added to the model one at a time and retained if their conditional TWAS association remained significant after Bonferroni correction for 247 tests. The same procedure was used to perform TWAS joint/conditional analysis of marginal SNP/QTL associations.

### Schizophrenia TWAS replication within PGC2

We used the individual-level PGC data^1^ to perform replication analyses and compare to summary-based results. Cis SNP-expression effect sizes were computed using the BSLMM software (as above) and predicted into each PGC sub-cohort to produce individual-level gene expression predictions. To evaluate the importance of SNP platform, two separate analyses were performed for the CMC, using all genotyped SNPs in the expression reference and all imputed HapMap3 SNPs, respectively. Association between predicted expression and SCZ was assessed using logistic regression with sub-cohort label and 10 principal components included as covariates. For the sub-sampling analysis, the full study was randomly split into discovery and replication. The association was then measured in discovery samples only, and any significant genes again tested for association in the replication samples. For sub-cohort analysis, one cohort was removed at a time (marked as replication) and the same process repeated.

### Gene-based risk scores

We adopted SNP-based polygenic risk score analysis^33^ to evaluate TWAS predictive accuracy and validation. Given a 1-by-*M* vector *z* of signed association statistics in the discovery study and an *N*-by-*M* matrix *X* of genetic values for the corresponding *M* genes in the replication study, we constructed a gene-based risk score *S* = *Xz*. The *M* genes were either all transcriptome-wide significant genes (gene risk score - GeRS) or all genes passing relaxed p-value thresholds (gene polygenic risk score - GePRS). This risk score was then tested against case/control status by a standard linear model *y* ~ *S* + *P* + *e* where *S* is the risk score and *P* is a matrix of principal components accounting for ancestry. The risk-score accuracy was then measured as the *R*^2^ from the above model less the *R*^2^ from the model *y* ~ *P* + *e* to account for ancestry, and converted to the liability scale assuming a prevalence of 1%.

For the TWAS using METSIM, YFS, and NTR expression reference panels, the cis-genetic component of expression was predicted in CMC samples. For the TWAS using the CMC expression panel, either the total expression was used (Figure S7) or the cis-genetic component of expression was estimated directly using BSLMM (equivalent up to a scaling factor to estimating genetic values by dropping each individual in turn). We stress that the case/control label from the CMC data was never used to identify the TWAS associations, and that the GeRS or GePRS from the CMC expression panel were thus evaluated against an independent CMC case/control phenotype. Ascertaining cases in the CMC expression panel may increase the frequency of causal variants and make the prediction more accurate than using a randomly ascertained expression panel, however, we observed little difference when performing the TWAS using an expression panel consisting of CMC controls only (Table S22).

### Power simulations for chromatin TWAS

Using real genotypes from the UK10K^58^ study, we simulated a model where SNPs X are causal for a chromatin peak (*C = X,β_X_* + *e_C_*) and chromatin is causal for expression (*E* = *Cβ_C_* + *e_E_*). 100 1MB loci were randomly selected across the genome, and causal SNPs in each locus were then randomly selected from common variants (MAF > 1%). Environmental noise was set such that SNPs explain 30% of the variance in a chromatin peak, and chromatin explains 30% of the variance in expression (consistent with our observations that expression 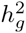 is ~ 10%). The TWAS was then performed using cis SNP-expression effect sizes computed by BSLMM either predicting the chromatin phenotype (with increasing sample size) into 1,000 independent individuals with expression or vice versa, and then performing an association between the predicted and measured phenotype. Separately, eQTL and cQTL association was computed for every common SNP in the locus using individuals with both expression and chromatin measured. To evaluate power at genome-wide significance, a TWAS association was reported as significant if it had *P* < 0.05 after correcting for (average 30 peaks per locus) × (20,000 genes) = 600,000 tests. For the QTL-based approach, given M SNPs in the locus, a SNP was reported as significant if it had an eQTL *P* < 0.05 after correcting for M × 20, 000 tests and a cQTL *P* < 0.05 after correcting for *M* × 30 × 20,000 tests. For a given chromatin sample size, the simulation was then performed at 100 random loci and 5 random seeds each, with the fraction of loci reported as significant by each method taken as the power (Figure S14). The TWAS simulation was separately performed under the null, using non-heritable chromatin and expression, and shown to be well-calibrated under the null (Figure s13).

### Individual-level chromatin TWAS

We used cis SNP-expression effect sizes computed by BSLMM scores in the four expression reference panels (including differentially spliced introns) to predict individual-level expression in the 45 CEU^18^ and 76^6^ YRI individuals with measured chromatin phenotypes. We retained only SNPs that were typed in both studies and removed strand-ambiguous SNPs. We did not perform any additional QC of the functional features, which were all previously PEER-adjusted and normalized^6,18^. We note that even though the YRI target samples are of different ethnicity, this prediction does not require an LD-reference panel and is therefore only expected to suffer loss in power (but not increased type I error) due to the differences in LD. For each predicted gene, we identified all chromatin peaks within a given window of the TSS (primary results used ±500kb) and tested each mark for association to predicted expression by linear regression.

### Multiple hypothesis correction for chromatin TWAS

The large number of correlated phenotypes analyzed - expression from five experiments and chromatin from nine experiments in two populations - allows for several approaches to multiple testing correction. For the chromatin TWAS, we corrected for the number of gene-peak pairs tested within a single expression reference and chromatin phenotype experiment (for example, number of gene-peak pairs when evaluating predicted CMC expression with the CEU:H3k27ac chromatin phenotype). This is directly comparable to the experiment-wide corrections applied in previous eQTL/cQTL analyses^6,18^. The same correction was applied for the SCZ/chromatin TWAS overlap: for example, the 44 SCZ TWAS genes identified using CMC expression were within 500kb of 1,528 total peaks in the CEU:H3k27ac experiment and “overlap” was reported for any peak that had a chromatin TWAS association *P* < 0.05/1, 528.

For comparison, we separately calculated the number of associations that were significant at 5% FDR across all molecular experiments. This yielded approximately 3.5× more chromatin TWAS associations and 1.2× more SCZ and chromatin TWAS associations (Table S2), demonstrating that the above experiment-wide Bonferroni correction strategy corresponds to a conservative study-wide FDR.

### eQTL/cQTL overlap analysis

We compared the chromatin TWAS to the traditional approach of identifying SNPs that are significant both as cQTLs and eQTLs in real data (Figure 4B). For each population and given distance to TSS, we performed this analysis in two stages. Stage 1: We identified all eQTLs that were significant after Bonferroni correction for the total number of SNP-gene pairs tested. When distance to TSS was the maximal allowed (500kb), this resulted in 355 eQTLs in the YRI and 579 eQTLs in the CEU data. Stage 2: From the set of significant eQTLs, we then looked for those that were also significantly associated with peaks from a given chromatin phenotype (for peaks within the given distance to TSS), after Bonferroni correction for the number of eQTL-peak pairs tested. In both stages the tests were only counted for the given chromatin phenotype (e.g. H3K27ac in CEU). This was compared to the chromatin TWAS analysis where each gene was tested against any peak within the given distance, and number of significant results reported after Bonferroni correction for total number of gene-peak pairs tested.

### Analysis of allele-specific expression

For each molecular phenotype (RNA-seq/ATAC-seq), we phased the locus using EAGLE2^59^ and evaluated haplotype allelic imbalance as follows. Given a “peak” SNP (for which there are RNA/ATAC-seq reads) and a “target” SNP (for which we want to evaluate allelic imbalance) we restricted to individuals that were heterozygous for both SNPs and counted the number of peak reads mapping to the REF/ALT haplotypes of the target SNP. We then assessed statistical significance for deviation from 50% balance using a binomial test. When the peak SNP and target SNP are identical, this is equivalent to a standard allelic-imbalance test across all heterozygous carriers in the peak. Duplicate reads were removed using samtools prior to all analyses and only variants with ≥ 10 reads at the peak SNP were tested. See Supplementary Materials for details on individual phenotypes analyzed.

### Ratio of cis-genetic covariance between chromatin-SCZ and expression-SCZ

To shed more light on the potential causal model, we sought to evaluate the evidence in support of two models of mediation: M_*CH*_, where SNP → chromatin → expression → disease; and M_*EX*_, where SNP → expression → chromatin → disease. Under the assumption of linear, additive variance across molecular phenotypes, this can be estimated via the ratio of genetic covariance (*cov_g_*) between chromatin-SCZ and expression-SCZ. The fraction of environmental variance on expression (*env_EX_*) under each model of mediation can then computed from the following equation (see Supp. Note for derivation):

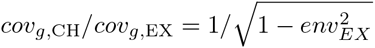

To compare these two models without bias from sample size or assay, we estimated the genetic covariance (*cov_g_*) using the PGC SCZ summary statistics and molecular data from the CEU/YRI individuals which had both chromatin and gene expression measured. For each SCZ/chromatin TWAS gene, we defined the target locus as ±50kb of the union of gene and peak boundary and estimated SCZ-chromatin and SCZ-expression *cov_g_* using cross-trait LD score regression^50^, restricting to the well-imputed HapMap3 SNPs and using in-sample LD. To estimate significance, background *cov_g_* was computed using the same procedure over 200 randomly gene-peak pairs within 500kb of the TSS. The observed and background estimates were then compared using the non-parametric Kolmogorov-Smirnov test. We note that for the YRI samples LD in the GWAS summary statistics is expected to differ from LD in the eQTL/cQTL data, and confound the raw estimate of *cov_g_*; however, because the random gene-peak pairs are also computed from the same population, we do not expect significance measured against this null to be inflated. We separately considered a partial correlation analysis, where each expression measurement was transformed to the residual of the associated chromatin peak in a standard linear regression (and likewise for each chromatin measurement). The two estimates of *cov_g_* were again computed from these partial phenotypes as described above. We caution that the estimate of e in the above equation was computed from an average across all loci, and could also be consistent with confounding from different levels of measurement error for ChIP-seq and RNA-seq, a mixture of models M_*CH*_ and M_*EX*_ that favors model M_*CH*_, or mediation by other unobserved molecular phenotypes.

